# The gene expression classifier ALLCatchR identifies B-precursor ALL subtypes and underlying developmental trajectories across age

**DOI:** 10.1101/2023.02.01.526553

**Authors:** Thomas Beder, Björn-Thore Hansen, Alina M. Hartmann, Johannes Zimmermann, Eric Amelunxen, Nadine Wolgast, Wencke Walter, Marketa Zaliova, Željko Antić, Philippe Chouvarine, Lorenz Bartsch, Malwine Barz, Miriam Bultmann, Johanna Horns, Sonja Bendig, Jan Kässens, Christoph Kaleta, Gunnar Cario, Martin Schrappe, Martin Neumann, Nicola Gökbuget, Anke Katharina Bergmann, Jan Trka, Claudia Haferlach, Monika Brüggemann, Claudia D. Baldus, Lorenz Bastian

## Abstract

Current classifications (WHO-HAEM5 / ICC) define up to 26 molecular B-cell precursor acute lymphoblastic leukemia (BCP-ALL) disease subtypes, which are defined by genomic driver aberrations and corresponding gene expression signatures. Identification of driver aberrations by RNA-Seq is well established, while systematic approaches for gene expression analysis are less advanced. Therefore, we developed ALLCatchR, a machine learning based classifier using RNA-Seq expression data to allocate BCP-ALL samples to 21 defined molecular subtypes. Trained on n=1,869 transcriptome profiles with established subtype definitions (4 cohorts; 55% pediatric / 45% adult), ALLCatchR allowed subtype allocation in 3 independent hold-out cohorts (n=1,018; 75% pediatric / 25% adult) with 95.7% accuracy (averaged sensitivity across subtypes: 91.1% / specificity: 99.8%). ‘High confidence predictions’ were achieved in 84.6% of samples with 99.7% accuracy. Only 1.2% of samples remained ‘unclassified’. ALLCatchR outperformed existing tools and identified novel candidates in previously unassigned samples. We established a novel RNA-Seq reference of human B-lymphopoiesis. Implementation in ALLCatchR enabled projection of BCP-ALL samples to this trajectory, which identified shared patterns of proximity of BCP-ALL subtypes to normal lymphopoiesis stages. ALLCatchR sustains RNA-Seq routine application in BCP-ALL diagnostics with systematic gene expression analysis for accurate subtype allocations and novel insights into underlying developmental trajectories.

## Introduction

Improved outcomes in B cell precursor acute lymphoblastic leukemia (BCP-ALL) – both, in pediatric and adult patients – have been achieved by precise risk stratification and target specific treatments. Molecular BCP-ALL subtypes and immunophenotype are the most important baseline prognosticators for BCP-ALL beside white blood cell counts and age. They inform risk-adapted treatments and targeted therapies. Currently, the revised WHO classification of lymphoid neoplasms, (WHO-HAEM5)^1^ and the International Consensus Classification of Myeloid Neoplasms and Acute Leukemia (ICC)^2^ have acknowledged 11 and 26 molecular defined BCP-ALL subtypes as distinct diagnostic entities, respectively, including 5 provisional entities (ICC classification). A total of 21 of these subtypes have been characterized by distinct gene expression profiles^3–8^, while the remaining subtypes^2,5^ are rare (IGH*::IL3*) or were defined by specific sets of underlying genomic drivers (Ph-like: ABL class / JAK-STAT / NOS) or their absence (*KMT2A*-/*ZNF384*-like). This heterogeneity of diagnostic subtypes exceeds the capabilities of cytogenetic (chromosome banding analysis, FISH) and molecular genetic methods (breakpoint specific PCR, MLPA, SNP-array / Array-CGH) which have so far been combined for identification of BCP-ALL subtypes. RNA-Seq enables identification of all BCP-ALL subtypes with a single method, establishing a new diagnostic standard. Further implementation as routine clinical diagnostic requires unified analysis methods. Calling of driver gene fusions^9,10^ is well established and novel approaches for the identification of hotspot single nucleotide^10^ variants and virtual karyoytpes^11^ exist. Yet only few approaches for systematic gene expression analysis are currently available.

Gene expression signatures represent the signaling equivalent of heterogeneous genomic driver alterations and have been used to define BCP-ALL subtypes. Initially, unsupervised clustering or prediction analysis for microarrays (PAM) were used to define subtype specific gene sets resulting in considerable heterogeneity regarding gene set definitions and subtype allocation of individual samples.^12^ More recent systematic approaches for BCP-ALL subtype allocations have employed machine learning methods to train classifiers for BCP-ALL subtype allocation mainly on pediatric ALL datasets.^13,14^ Yet the optimal method still needs to be defined – especially for rare and difficult to classify subtypes and subtypes with predominance in adults. Additionally, correct assignment of samples, which do not fall into established subtype categories either due to interfering biological conditions (e.g., low blast count, poor RNA quality) or because these samples represent novel candidate subtypes, remains a challenge. In addition to molecular subtype definitions, gene expression profiles might be informative for clinical baseline parameters such as leukemic blast proportion, immunophenotype or more detailed analysis of lymphopoiesis trajectories underlying BCP-ALL development. However, systematic approaches and especially RNA-Seq data that link BCP-ALL subtypes to human B lymphopoiesis differentiation stages are lacking.

Here we describe ALLCatchR, a machine learning based classifier pretrained for allocation of BCP-ALL gene expression profiles to all 21 gene expression defined molecular subtypes of WHO-HAEM5 and ICC classifications. High accuracies in independent validation cohorts are achieved by integrating machine learning and gene set based nearest neighbor models into a compound classifier. ALLCatchR infers clinical baseline variables such as blast proportion and patient’s sex from RNA-Seq data and provides a putative differentiation stage of origin based on our newly established reference of human B lymphopoiesis. ALLCatchR sustains routine diagnostic application of RNA-Seq with systematic gene expression analysis providing subtype allocations and insights into underlying biology for further exploratory analysis.

## Material/Subjects and Methods

### Aggregation of a 3,532 sample BCP-ALL transcriptome reference data set

To establish a classifier for BCP-ALL molecular subtype allocation, we aggregated RNA-Seq count data from n= 3,532 BCP-ALL patients including 64.5% pediatric^5–7,13^ and 35.5% adult^3– 5,8,13^ cases combined from 6 independent datasets (**Figure 1A**; **Supplementary Table S1**). Excluded were samples with multiple subtype assignments (n=116), multiple representations of the same patient (n=44), subtypes which are not part of WHO-HAEM5 / ICC classification (Low hyperdiploid, *IDH1/2*; n=55) or which are mainly defined by absence of a genomic driver (*KMT2A*-like, *ZNF384*-like; n=9). Molecular BCP-ALL subtype allocations were performed for n=2,887 samples in the original studies based on genomic drivers and corresponding gene expression signatures. Subtype-defining genomic events were identified in >90% of cases either by RNA-Seq (gene fusions, hotspot single nucleotide variants, virtual karyotypes) or by genomic profiling (whole genome- / whole exome- / gene panel sequencing, SNP-arrays, array-CGH). A total of n=421 samples where defined ‘unassigned’ or ‘B-other’ in the original studies. All BCP-ALL molecular subtypes from current WHO-HAEM5 or ICC classifications which were characterized by distinct gene expression signatures in their original description (n=21) were represented in the data set (not included: IGH:*:IL3, KMT2A*-like, *ZNF384*-like. Ph-like was considered one subtype without sub-division. CEBP/ZEB2 subtype lacks final definitions so far and was defined here as ‘CEBP’ by presence of *IGH::CEBPA/CEBPE/CEBPD* fusions and absence of other drivers.) (**Supplementary Table S2**). Raw read counts for 15,728 protein-coding genes represented in all cohorts were used including heterogenous sequencing approaches (poly-A selection / depletion of ribosomal RNAs), sequencing depths and different read count quantification methods before normalization (*log10(count + 1)*, followed by z-transformation and scaling between 0-1).The data set was split into a data set used for training of the classifier (n=1,869) and 3 hold-out studies (n=1,018) for independent validation both representing all analyzed BCP-ALL subtypes (**Figure 1A, Supplementary Figure S1**).

**Figure 1.**
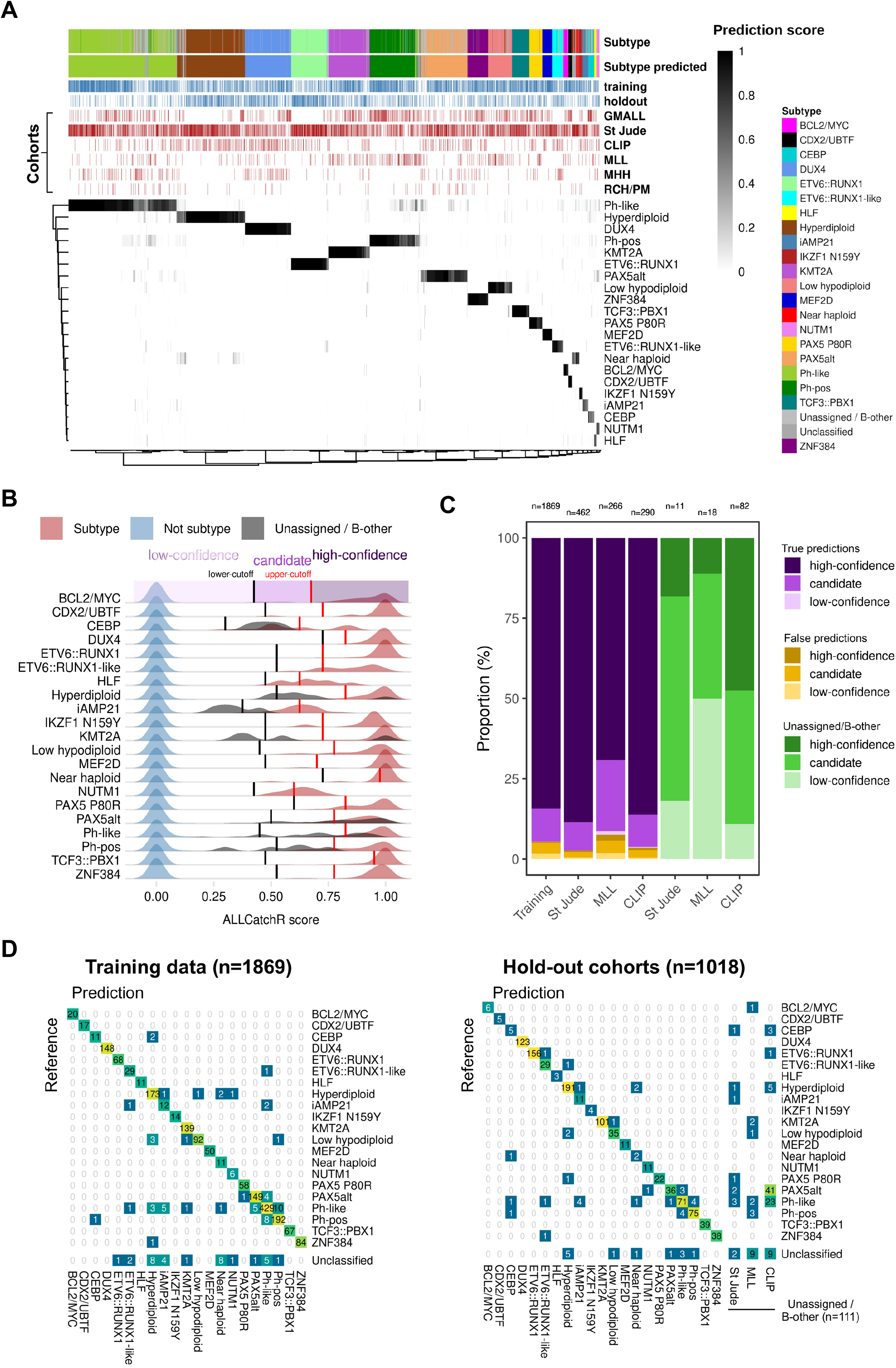
ALLCatchR predicts molecular BCP-ALL subtypes based on gene expression count data with high accuracy. **(A)** Heatmap showing the prediction scores for 21 gene expression defined BCP-molecular subtypes (WHO-HAEM5 / ICC) in n=3,308 samples of the entire BCP-ALL cohort (after removal of duplicate samples and samples with two primary subtype allocations; n=217) samples. Molecular subtypes had been defined in the six original studies (GMALL, St Jude, CLIP, MLL, MHH and RCH/PM) based on genomic driver aberrations and corresponding gene expression signatures in n=2,887 cases (ground truth). Remaining cases were deemed ‘unassigned’ or ‘B-other’. ALLCatchR scores are shown for the combined data set of training and hold-out cohorts. **(B)** Cutoffs were defined for each BCP-ALL subtype based on distribution of all ALLCatchR scores in every subtype. Cutoffs for ‘high confidence predictions’ were defined to include >90% of all samples allocated to these subtypes in the original data set, resulting in 0.989 accuracy of these predictions in independent hold-out data set. Cutoffs for ‘Candidate predictions’ were defined to reliably exclude samples from other subtypes, providing a reliable orientation for further validation of subtype assignment based on genomic drivers (accuracy: 0.851). ‘Low-confidence’ predictions indicate samples from different subtypes or samples where subtype allocation cannot be performed. These were considered ‘unclassified’ for further analysis. **(C)** The proportions of confidence categories for true and false predictions in the training and hold-out data sets are shown. A prediction was considered ‘true’ if the sample received the same subtype allocation as in the original study. ‘False’ predictions represent allocations to other subtypes than the subtype assigned in the original study. For comparison, ‘unassigned’ / ‘B-other’ samples from the holdout data sets are shown. **(D)** Confusion matrices relate ALLCatchR predictions to the ground truth in training samples (left) and holdout cohorts (right). By design, the training cohort did not contain ‘unassigned’ / ‘B-other’ samples. In the hold-out data, n=111 samples had been defined as ‘unassigned’ / ‘B-other’ and predictions for these are also shown. Supplementary Figure S5A and Supplementary Table S5 indicate how ALLCatchR predictions in ‘unassigned / B-other’ samples are supported by corresponding genomic drivers in 72.1% of ‘high confidence’ and 27.1% of ‘candidate’ predictions.

### Integration of machine learning and gene set based nearest-neighbor models for BCP-ALL subtype allocation

To perform molecular subtype allocation based exclusively on gene expression data, we developed ALLCatchR, a classifier which integrates linear support vector machine (SMV) and nearest-neighbor association models for BCP-ALL subtypes derived from the training data (**Supplementary Figure S1**) Feature selection (LASSO)^15^ using the glmnet package^16^ was used to extract BCP-ALL subtype defining gene sets, resulting in 2,802 genes with high discriminative power for 21 molecular subtypes (**Supplementary Figure S2, Supplementary Table S3**). First, we used this gene set to train five different machine learning classifiers using two feature selection methods^15,17^ of which linear SVM^18^ performed best independent of the feature selection method used (**Supplementary Figure S3**). This resulted in a high accuracy (0.963) of subtype prediction in the training data. However, linear SVM is restricted to predefined classes and does not compute probabilities for individual subtype predictions, which prevents it from correctly handling cases which are unassigned or ambiguous due to multiple drivers or which represent novel candidates. To achieve a probabilistic compound model, we incorporated single sample gene set enrichment analyses (ssGSEA) using singscore^19^ of the same subtype-defining LASSO gene sets. By this approach, batch effects between cohorts were removed (**Supplementary Figure S4**) Euclidean distance of each test sample to each training sample was computed and the 10 nearest neighbors were considered for subtype allocations of each test sample (accuracy for subtype prediction based on highest enrichment for each sample: 0.912). Both models - SVM linear predictions and sample-to-samples-distances in subtype-defining gene sets – were integrated into our newly established compound classifier, ALLCatchR, which provides dynamic ranges of subtype-specific probability scores (**Figure 1A**). To achieve a better separation between highly similar high hyperdiploid and near haploid ALL, both subtypes where first represented as one class in the overall classifier (NH/HeH) and then separated by a second 2-class compound classifier with a similar design as the overall classifier.

### Development of an RNA-Seq reference of human B-lymphopoiesis

Bone marrow samples from healthy adult donors (n=4, M:F=1:3, age: 27-39 years, study registration DRKS00023583, ethical approval of ethics committee, Kiel University: D 583/20) were subjected to immunodensity cell separation (RosetteSep, STEMCELL Technologies; Inc., Vancouver, BC, Canada; purging: CD16, CD36, CD66b, CD235a, CD3). Non-depleted cells were stained with a 9-color antibody panel (**Supplementary Table S4**) and FACS-sorted (FACSAria™ fusion; BD Biosciences, Franklin Lakes, NJ, USA) to 7 lymphoid differentiation stages. RNA was extracted from 5,000-320,000 cells per differentiation stage (AllPrep™ DNA/RNA Micro Kit, Qiagen, Venlo, Netherlands) and subjected to ultra-low-input RNA sequencing after generation of stranded sequencing libraries (SMART-Seq® Stranded Kit, Takara Bio Inc., Kusatsu, Shiga, Japan; NovaSeq 6000, Illumina, San Diego, CA, USA).

## Results

### ALLCatchR performs BCP-ALL molecular subtype allocation with high accuracy

We used aggregated BCP-ALL gene expression profiles (n=3,532 samples, n=6 cohorts) to develop ALLCatchR, a pre-trained machine learning classifier which performs BCP-ALL molecular subtype allocation based on gene expression alone (detailed in ‘Methods’). ALLCatchR provides probability scores for each sample and all gene expression defined BCP-ALL subtypes (**Figure 1A**). Unsupervised clustering of ALLCatchR scores groups samples according to subtype across cohorts and age groups. For final subtype allocation, we defined subtype-specific cutoffs based on the comparison of probability scores from samples belonging to the corresponding subtype and all remaining samples of the cohort (**Figure 1B**). This resulted in 1.) high-confidence predictions, 2.) candidate predictions and 3.) low-confidence predictions i.e., unclassified samples. Cutoffs for ‘high-confidence’ predictions were defined to include >90% of correct predictions. Cutoffs for candidate predictions were defined to exclude all samples from other subtypes but allowed unassigned/B-other samples (n=111; **Figure 1B**). In the training data, 84.6% of samples achieved high confidence predictions with an accuracy of 0.997, while 13.7% achieved candidate predictions with an accuracy of 0.797 to guide further validation based on genomic drivers in well pre-specified directions (**Figure 1C**). Only 1.7% of samples achieved low-confidence predictions and were considered ‘unclassified’. To validate ALLCatchR performance, we used independent validation data from 3 hold-out cohorts (n=1018; **Supplementary Figure S1A**), not previously seen by the classifier. A total of n=1006 (98.8%) samples was allocated to one of 21 subtypes (high-confidence and candidate predictions) with an accuracy of 0.957, demonstrating the feasibility of highly accurate subtype allocations based on gene expression alone. ‘High-confidence’ and ‘candidate’ predictions were achieved in 83.7% and 15.1% of samples with accuracies of 0.989 and 0.851 respectively. A total of n=32 samples (3.1%) were assigned to the wrong subtype or received no subtype allocation (n=12; 1.2%). Most prominent misclassifications affected Ph-like-to Ph-pos predictions or vice versa (n=8) or subtype allocations of aneuploid subtypes (n=20), **Figure 1D**). The majority of misclassified samples (n=23/33; 67.6%) had received candidate predictions, supporting the need to validate these predictions based on genomic drivers.

### ALLCatchR provides subtype allocations for previously ‘unassigned / B-other’ samples

In addition to the n=1018 hold-out samples with assigned subtype, n=111 samples had been defined as ‘unassigned / B-other’ (n=107) or were identified as ‘non-Ph-like *CRLF2*-rearranged’ (n=4) in the original studies (**Figure 1C, D**). ALLCatchR concordantly identified n=20 (18.0%) of these as ‘unclassified’ (**Figure 1D, Supplementary Figure S5**). However, n=43 (38.7 %) and n=48 (43.2 %) cases received ‘high-confidence’ or ‘candidate’ predictions respectively (**Figure 1D**). Analysis of available RNA-Seq gene fusion calls or cytogenetic profiles and/or virtual karyotyping (WGS / SNP-arrays) identified driver candidates supporting the corresponding subtype allocations in n=31 (72.1%) of ‘high-confidence’ and n=13 (27.1%) of ‘candidate’ predictions (**Supplementary Table S5; Supplementary Figure S5**). These newly suggested subtype allocations consisted of PAX5alt predictions (n=25) which had not shown a clear PAX5alt gene expression profile in the original cohort (n=1), or which were contributed from the CLIP cohort where this subtype had not been annotated previously. Next, n=11 *CRLF2*-rearranged cases from CLIP and St Jude cohorts without Ph-like gene expression profile in the original cohorts received ALLCatchR Ph-like predictions. Among the remaining n=7 samples, one case with an ALLCatchR high-confidence KMT2A prediction was found to harbor a *KMT2A* partial tandem duplication by WGS (**Supplementary Figure S5**). To the best of our knowledge, this is the first identification in BCP-ALL of this aberration which is recurrently observed in acute myeloid leukemia. In a second of these n=7 cases, an IGH:*:MYC* gene fusion was identified in support of a *BCL2/MYC* ALLCatchR prediction. Further ALLCatchR high-confidence predictions for ‘unassigned / B-other samples’ without corresponding drivers included PAX5alt (n=9) and Ph-like (n=3) predictions, which generally are defined in a proportion of samples by gene expression alone. Thus, ALLCatchR suggested molecular subtype allocations in previously ‘unassigned’ cases with atypical and less well-defined gene expression signatures and supported the identification of novel driver candidates.

### High accuracy of ALLCatchR predictions is observed across cohorts and molecular subtypes

The accuracy of predictions was consistently high in the training and hold-out data, with 0.952 and 0.957, respectively. Almost congruent predictions were achieved in St Jude and CLIP cohorts with accuracies of 0.978 and 0.965, respectively. In the MLL hold out set the accuracy was slightly lower with 0.914 (**Figure 2A**). Of note, the MLL cohort includes real-world adult BCP-ALL data from a diagnostic laboratory with more permissive pre-selection cutoffs (e.g., blast counts) indicating that ALLCatchR achieves reliable predictions also in less pre-selected samples. Despite the overall high accuracies, classification performance varied between molecular subtypes (**Figure 2B**). ALLCatchR achieved specificities >0.99 for all 21 subtypes, both in training and testing data sets. The average sensitivity across subtypes was 0.919±0.145 and 0.911±0.167 in the training and hold-out data, respectively. For n=17/21 subtypes, sensitivities were ≥0.85 both on training and hold-out data, together including n=2,781 patients (96.3%; **Figure 2B**). Only 4 remaining subtypes (n=106 samples, 3.7% of entire cohort) achieved sensitivities below 0.85 (NUTM1, *CEBP*, iAMP21 and Near haploid) which was mainly related to the small number of samples representing these subtypes, limiting both.

**Figure 2.**
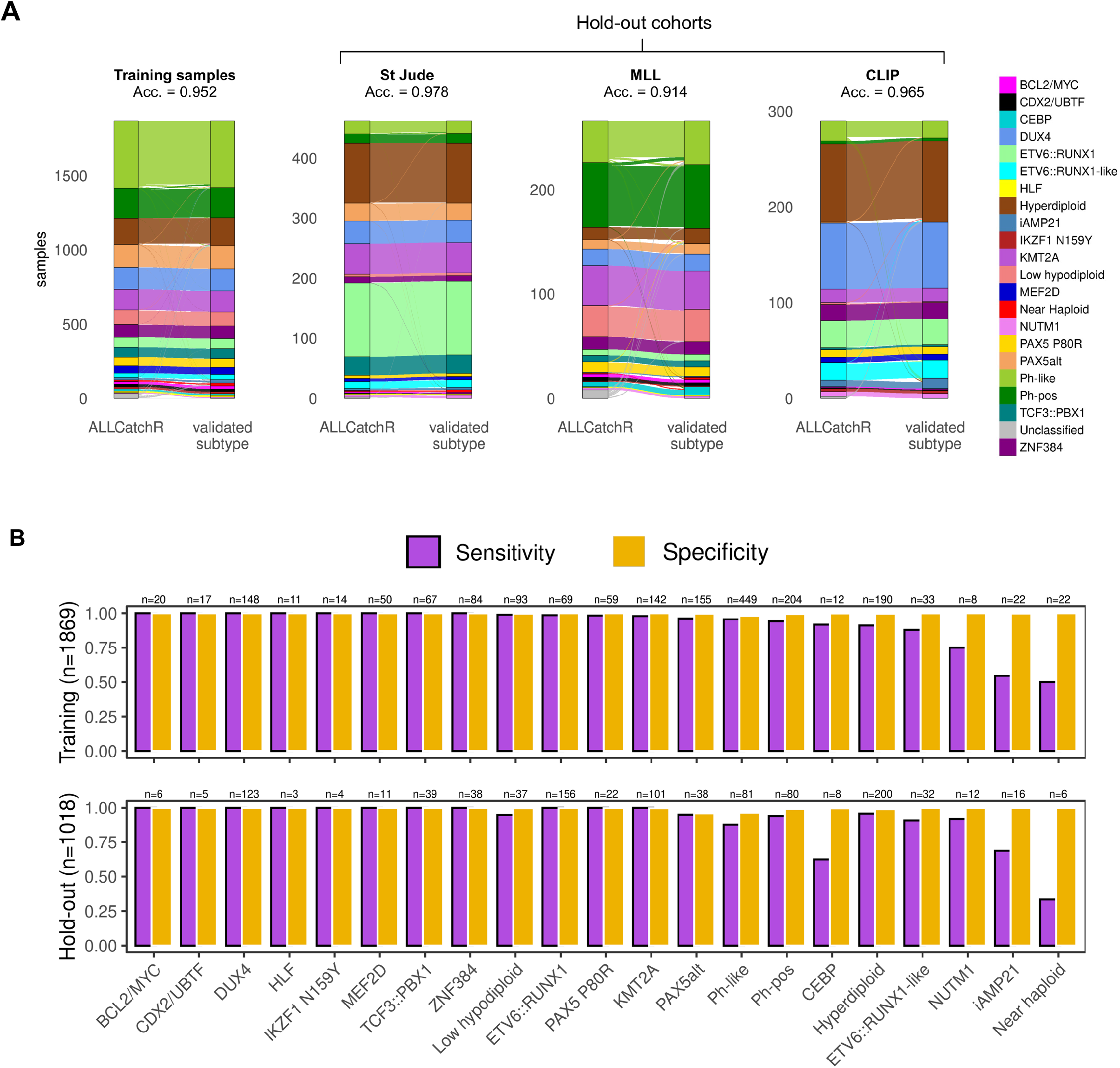
ALLCatchR accuracy for subtype allocation is consistently high across cohorts and BCP-ALL subtypes. **(A)** Sankey diagrams indicate ALLCatchR subtype allocations and corresponding subtype validated ground truth in the training cohort and the individual holdout data sets. ‘Acc.’ Indicated accuracy in the corresponding data set. **(B)** Bar charts indicated sensitivity and specificity for the individual subtypes in the training and hold-out data. Validated ground truth was used to define true positive cases, i.e. belonging to this subtype and true negative cases, i.e. not belonging to this subtype. Values were obtained as fraction of true positive cases from all cases defined by ALLCatchR as belonging to this subtype (sensitivity) and as fraction of true negative cases from all cases defined by ALLCatchR as not belonging to this subtype (specificity).

### ALLCatchR subtype allocation outperforms current tools

Recently, two tools - ALLSorts^13^ and Allspice^14^ - were independently developed for BCP-ALL subtype allocation based on gene expression profiles. In comparison to these, ALLCatchR provides comprehensive subtype-allocation to all gene expression defined WHO / ICC subtypes (n=21), including *CEBP* and *CDX2/UBTF*, which are missed by both tools. For performance comparison, n=2,887 samples with established subtype definitions (ALLCatchR training and validation data sets) were predicted with ALLSorts and Allspice (**Supplementary Figure S6**). ALLSorts performed well with an accuracy of 0.913 but left more samples ‘unclassified’ (n=145), compared to ALLCatchR (n=44). The largest difference was observed in the MLL holdout data, where ALLSorts achieved an accuracy of 0.771 compared to 0.914 accuracy for ALLCatchR (**Supplementary Figure S6**). An inferior performance in ALLSorts was mainly related to missed sample classifications (ALLSorts ‘unassigned’: n=43 (16.17%); ALLCatchR ‘unassigned’: n=8 (3.01%)). The MLL data set represents real-world data from a diagnostic laboratory with less stringent pre-selection of samples and thus represent a *bona fide* challenge for the tools. Allspice leaves more samples of the same cohort (n=2,887) unclassified resulting in accuracies of 0.629 in the training and 0.719 in the hold-out studies. However, for samples that could be assigned to a subtype by Allspice, prediction performances were comparable to ALLCatchR (**Supplementary Figure S6**). In summary, ALLCatchR achieves a higher accuracy for molecular subtype predictions, assigning more samples to the correct subtype including all gene expression defined subtypes.

### Gene expression-based modelling predicts clinical baseline variables

Blast count proportions impact accuracy of gene expression based molecular subtype allocation, as sequencing reads from non-leukemic compartments contribute to bulk transcriptome profiles. To infer sample blast proportions, we trained two machine learning regression models on data sets of our combined cohort with available blast counts obtained by manual counting or flow cytometry (GMALL, MLL) and used these as well as the RCH/PM cohort for validation. Blast count predictions from single cohorts achieved good accuracies when applied to each other (**Figure 3A-B**) with a high concordance between USKH and MLL training sets (**Figure 3B**) which were therefore combined for the final classifier. Only 1.85% of samples with high confidence subtype predictions had blast count predictions <50% while these were observed in 9.83% of candidate predictions and in 17.95% of unclassified samples of the entire cohort (**Supplementary Figure S7**). Thus, ALLCatchR can identify a subset of samples with worse performance for subtype allocation due to lower blast infiltration. Gene expression profiles were also informative for patient’s sex and disease immunophenotype. To enable gene expression based cross-validation of these important clinical baseline characteristics, we implemented sub-classifiers to the samples immunophenotype (pro-B vs. common-/pre-B ALL; accuracy of 0.871 in the validation data) and patient’s sex (accuracy: 0.991 in validation data set, **Figure 3C**). ALLCatchR thus provides a cross-validation of clinical baseline variables and allows imputation of missing values.

**Figure 3.**
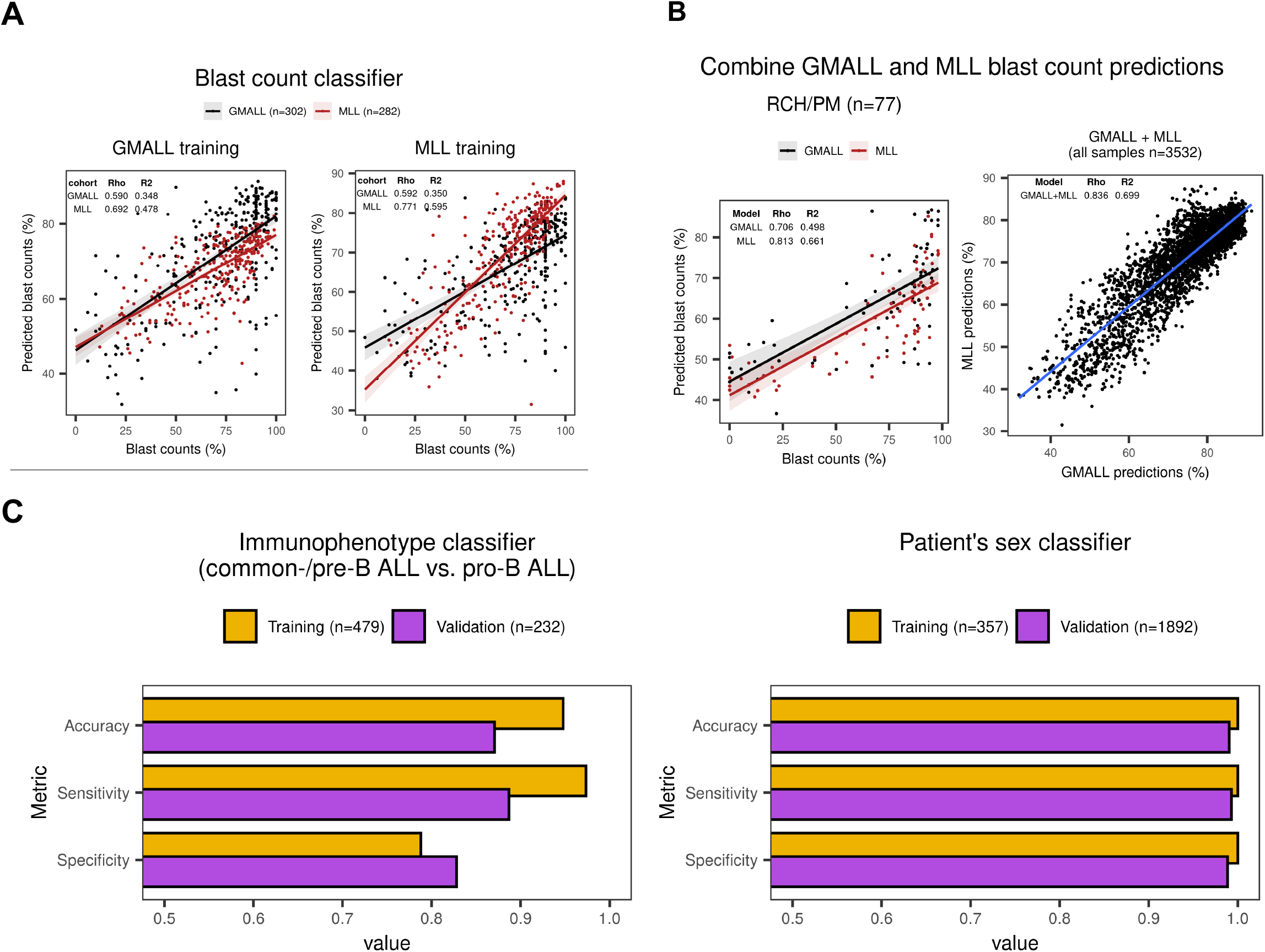
ALLCatchR predicts sample blast counts, patient’s sex and immunophenotype based on gene expression data. **(A)** For GMALL (n=302), MLL (n=282) and RCH/PM (n=77) sample blast counts obtained by cytology or flow cytometry were available. GMALL and MLL cohorts were separately used for training two classifiers in a 10-fold cross-validation scheme with the same machine learning algorithms used for subtype prediction. GMALL and MLL classifiers were validated on each other, and both were validated on the RCH/PM data. Best performing methods in terms of the Root Mean Squared Error (RSME) on the training data are shown. Training two classifiers on independent data sets allowed for the validation on each other and both were combined for final predictions. Blast count predictions had a good correlation to measured counts i.e., rho=0.590 in GMALL and rho=0.771 in MLL. Moreover, predicting MLL samples with the classifier trained on GMALL achieved a similar performance as the classifier trained on MLL samples and *vice versa*. **(B)** Since both, GMALL and MLL classifiers had a good performance and were generalizable, predictions from both are combined in ALLCatchR. **(C)** Sub-classifiers for immunophenotype and patient’s sex were developed using SVMlinear and ranger machine learning models respectively. An immunophenotype classifier was trained on GMALL samples (n=413 common-B / pre-B and n=66 pro-B) and validated on MLL data (n=168 common-B / pre-B and n=64 pro-B) with available EGIL immunophenotypes. A patient sex classifier was trained on n=357 GMALL samples (female=165, male=192) analogous to the subtype classifier. For validation n=1892 St Jude samples with known sex (female=850, male=1042) were used. Corresponding accuracies, sensitivities and specificities are shown for these sub-classifiers.

### Shared gene expression patterns suggest distinct cells of origin for BCP-ALL subtypes

The cell of origin for BCP-ALL cases remains to be defined, with immunophenotyping according to EGIL criteria^20^ representing a framework for orientation. An improved understanding of underlying lymphopoiesis trajectories is especially warranted regarding current immunotherapies which rely on differentiation-stage- and lineage-specific markers as therapeutic targets. To map BCP-ALL subtypes to underlying B lymphopoiesis trajectories, we established a reference of normal human B lymphopoiesis for 7 differentiation stages from hematopoietic stem cells to mature bone marrow B cell subsets (**Figure 4A**), based on established definitions^21^. Expression profiles were obtained from ultra-low input RNA-Seq of FACS sorted bone marrow samples of healthy adult donors (n=4). Unsupervised analysis of variable expressed genes grouped samples according to the developmental course (**Figure 4B**). Stage specific gene sets were obtained by multi-comparison ANOVA on normalized counts (vst), yielding well discriminative definitions (**Figure 4C**; **Supplementary Table S6**). Analysis of immunoglobulin rearrangements using droplet PCR indicated initiation of D_H_-J_H_ rearrangements in sorted pro-B cells while V_H_-(D)J_H_ rearrangements were first observed in pre-B II Large cells and class switch recombination occurred exclusively in the most mature B cells, providing an immunogenomic differentiation trajectory^22^ which independently confirms our sorting strategy (**Supplementary Figure S8**). We implemented this newly established model of human B lymphopoiesis in ALLCatchR using ssGSEA to define the proximity of each BCP-ALL sample to all 7 lymphopoiesis stages (**Figure 4D; Supplementary Figure S9**). Medians of theses enrichment scores across samples revealed distinct patterns of enrichments suggesting shared stages of origin for BCP-ALL subtypes (pro-B / pre-B I / pre-B I to pre-B II Large transition / pre-B II Large; **Supplementary Figure S9**) with similar patterns in pediatric and adult data sets (**Supplementary Figure S10**). Most BCP-ALL subtypes and the majority of all cases showed highest similarity to the pre-B I stage (Figure 4D). *KMT2A*-rearranged and *PAX5* P80R ALL showed a clearly distinct enrichment pattern favoring an earlier pro-B differentiation stage of origin (**Figure 4E**). In contrast, *CEBP, HLF, IKZFN1 N159Y, MEF2D, NUTM1* and *TCF3::PBX1* were grouped in a cluster with highest enrichment in transition of pre-B-I to pre-B-II large stage and *BCL2/MYC* showed the highest degree of similarity exclusively to pre-B II Large differentiation stage (**Figure 4D**). These observations confirm expectations for the extremes of this trajectory (*KMT2A* and *BCL2/MYC*).^23,24^ A recently reported mouse model of *PAX5* P80R ALL^25^ established a pro-B differentiation arrest as initial event in *PAX5* P80R homozygous models, supporting a pro-B origin of this leukemia subtype or at least an altered PAX5 function inducing a pro-B like phenotype in P80R mutated cases (**Figure 4E**). Thus, specific enrichment patterns of normal lymphopoiesis are shared between molecular subtypes, suggesting distinct stages of transition from normal to leukemic lymphopoiesis. We have included this model in ALLCatchR. Comparison of EGIL immunophenotypes to gene-expression-defined stages of origin indicated expected enrichments (pro-B stage in pro-B immunophenotype / pre-B II Large in pre-B immunophenotypes; (**Figure 4F**) but nearly all gene-expression-based differentiation stages were represented in each immunophenotype. BCP-ALL subtypes were more closely related to gene-expression-based differentiation stages as to EGIL immunophenotypes, suggesting that ALLCatchR identifies developmental underpinnings of BCP-ALL drivers at higher resolution.

**Figure 4.**
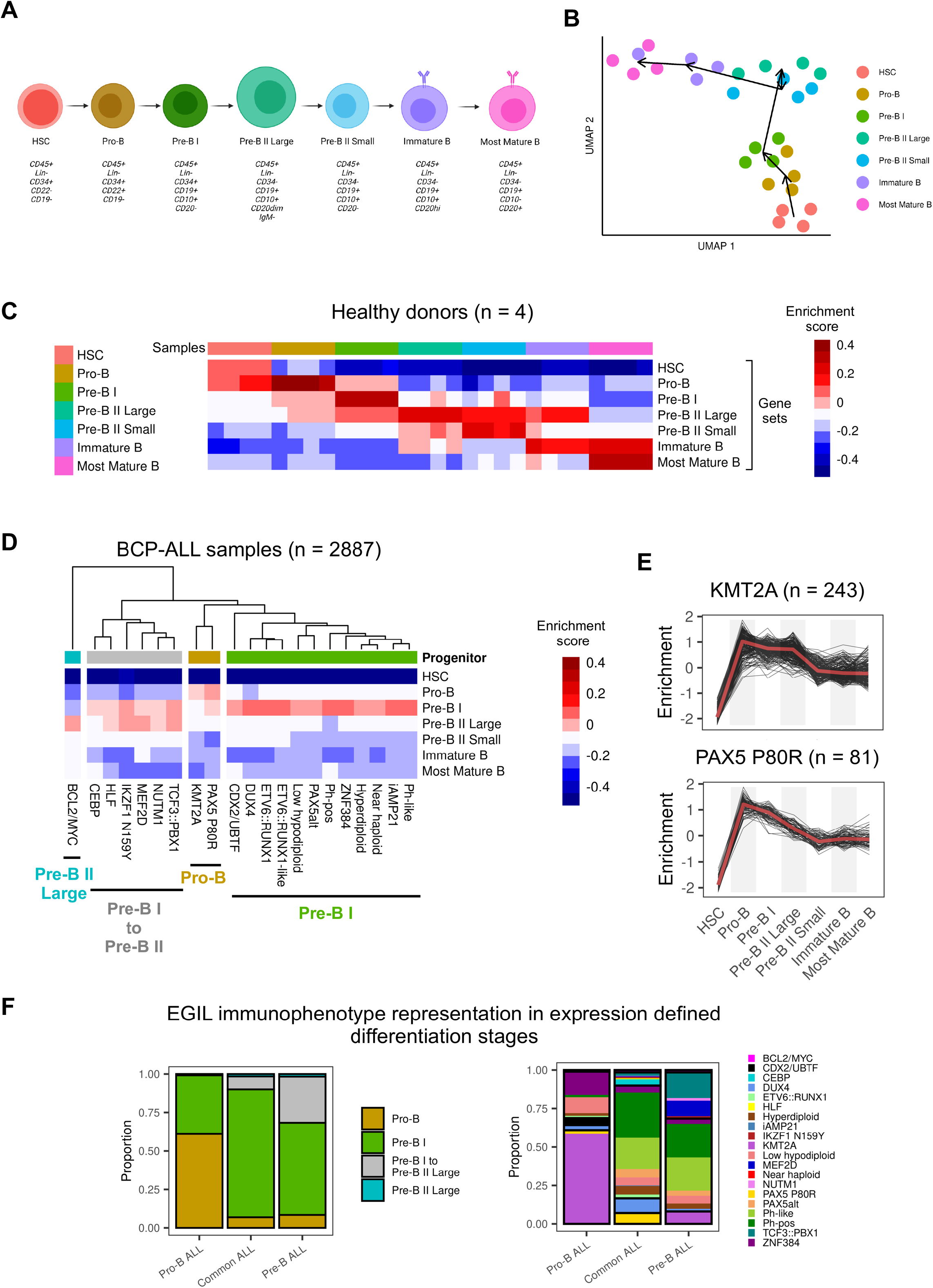
ALLCatchR identifies B cell developmental trajectories underlying BCP-ALL subtypes. **(A)** To establish a reference map of human B lymphopoiesis, we obtained bone marrow samples from healthy adult donors (n=4) and used a 9-color antibody panel for FACS sorting of 7 B lymphopoiesis stages following described definitions^21^ after pre-enrichment of wanted populations. Lin-selection included CD3, CD33, CD56, CD14, CD66c, CD138. Antibodies used are shown in Supplementary Table S4. Supplementary Figure S8 shows immunogenomic profiling of immune gene rearrangements in support of the applied sorting strategy. **(B)** Ultra-low input RNA-Seq was performed for total RNA to obtain stage-specific gene expression. Uniform manifold approximation plot (UMAP) shows clustering of human B lymphopoiesis stages based on 400 most variable expressed genes. **(C)** Multi comparison ANOVA on normalized (vst) count data was performed to obtain differentiation-stage specific gene sets. Heatmap depicts single sample gene set enrichment analyses (singscore)^19^ of B lymphopoiesis subsets (columns) to stage defining gene sets (rows). **(D)** BCP-ALL samples with known subtype allocation (n=2,887) were used for single sample gene set enrichment analysis with B lymphopoiesis-specific gene sets obtained from (C). Supplementary Figure S9 shows enrichment patterns of individual samples from all BCP-ALL subtypes for all differentiation stages. Heatmap depicts averaged enrichment scores for all BCP-ALL subtypes and all B lymphopoiesis stages grouped by unsupervised clustering. Normal progenitors with closest proximity to BCP-ALL subtypes representing putative cells-of-origin are annotated on top. Supplementary Figure S10 provides separate analyses for pediatric and adult patients indicating a high degree of similarity. (**E**) *KMT2A* rearranged and PAX5 P80R ALL had both the highest enrichment towards pro-B supporting a shared developmental origin (also depicted in Supplementary Figure S9). (**F**) Comparison of gene expression defined differentiation stages and EGIL immunophenotypes are shown for n=711 samples with available gene expression data.

### BCP-ALL subtype-defining gene sets indicate shared signaling trajectories

Definitions of BCP-ALL subtype specific gene expression signatures depend on the size and composition of the remaining cohort used as comparator. We made use of the aggregated transcriptome profiles of 21 BCP-ALL subtypes to define subtype specific gene expression profiles based on the largest data set (n=3,532) available till date, representing different age groups, cohorts, and sequencing methods. UMAP clustering of all samples according to LASSO selected subtype specific gene sets indicated a clear separation of molecular subtypes independently of the contributing cohorts (**Figure 5A**). To characterize subtype specific gene expression profiles beyond top discriminative features, we performed differential gene expression analysis for each subtype compared to the remaining cohort. A median of 673 differentially expressed genes per subtype were identified (range: 144– 1465; fold change: <1.5-log2-fold, FDR: <0.001; **Figure 5B**). Overlap between these gene sets was very low (**Supplementary Figure S11**) indicating that subtype-specific differences are represented in broad gene regulatory programs. Subtype specific gene expression profiles were provided as a resource in **Supplementary Tables S7-28**. To explore the potential of this dataset to reveal underlying biological functions, we performed ssGSEA for canonical signaling pathways (MSigDB Hallmark / KEGG gene sets). Analysis of pathways top differentially enriched in BCP-ALL subtypes (one-way ANOVA) indicated previously unrecognized clusters of subtypes with enrichment in cytokine receptor / JAK-STAT signaling (Ph-pos, Ph-like, *ZNF384*, Hyperdiploid, iAMP21) or WNT-/beta catenin/ hedgehog signaling (*ETV6::RUNX1* and -like, *CDX2/UBTF*), which together represented the majority of subtypes with a putative pre-B-I cell of origin (**Figure 5C**). For the remaining subtypes an enrichment in MYC-/MTOR signaling was observed in subtypes of both, a more and less mature differentiation stage of origin (pro-B: *KMT2A*, PAX P80R / pre-B I to pre-B II large: *BCL2/MYC, IKZF1 N159Y, MEF2D*; **Figure 5C**). Thus, enrichment analysis for canonical signaling pathways independently grouped together BCP-ALL subtypes form similar underlying B lymphopoiesis differentiation stages. ALLCatchR not only provides a systematic gene expression analysis for accurate identification of molecular BCP-ALL subtypes but also enables insights into underlying disease biology which is closely interconnected with subtype nosology.

**Figure 5.**
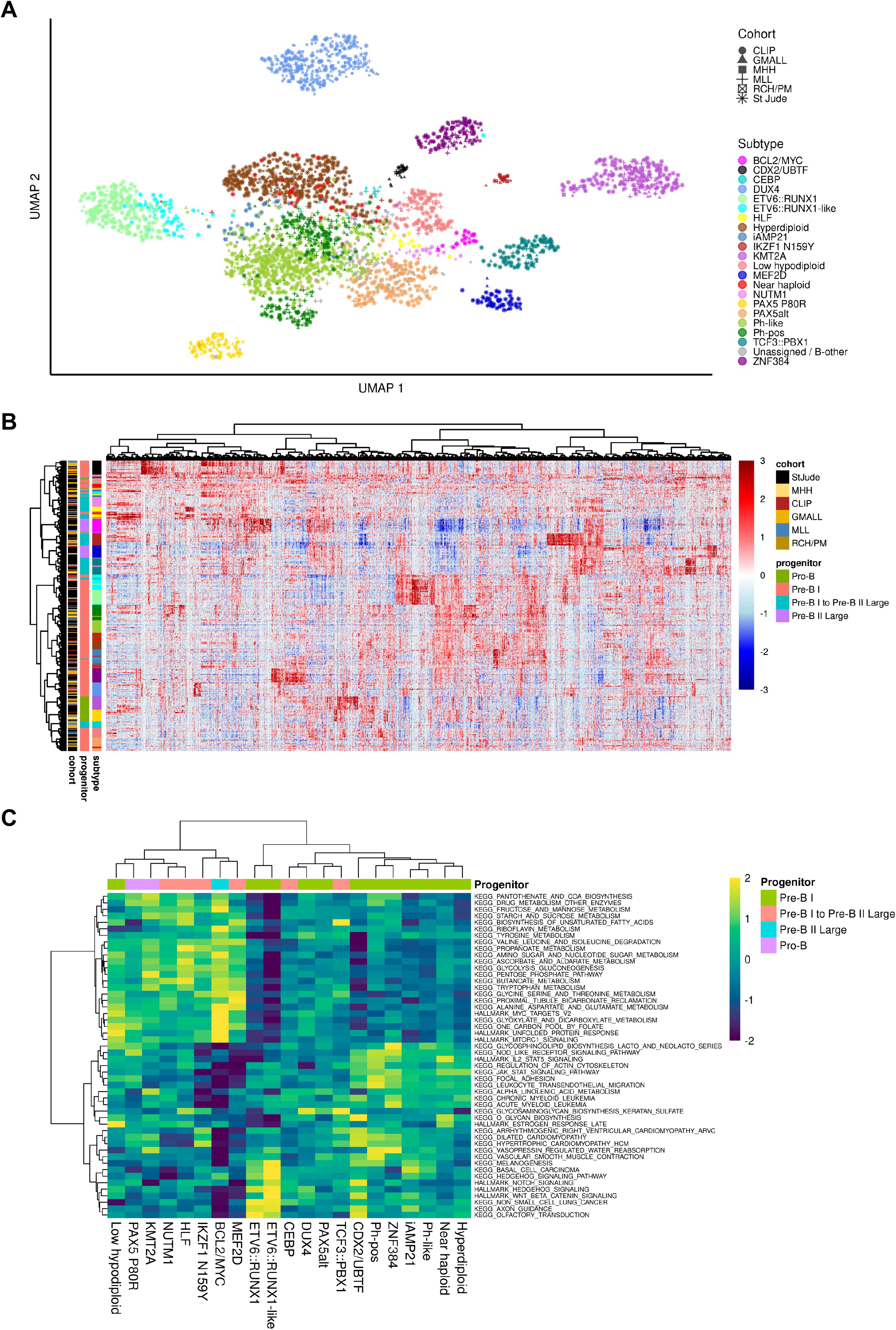
The gene expression landscape in BCP-ALL. **(A)** UMAP plot showing all n=3,308 samples used in this study. Count data from the six data sets was batch corrected using the sva package^33^ and TPM values calculated. The plot is based on 2,802 genes selected by LASSO for training of ALLCatchR. Cohorts are highlighted as shape. **(B)**. ALLCatchR predictions were used to define samples which best represented their respective molecular subtype. A total of n=20 top ranking samples per subtype (exceptions with lesser samples available: HLF n=14, *CEBP* n=16, *NUTM1* n=17, *IKZF1 N159Y* n=18) were used to obtain a homogenous data set representing all 21 BCP-ALL subtypes (n=405). Differential gene expression analyses for each subtype versus the remaining cohort using DESeq2^34^ revealed 5,110 differentially expressed genes (cutoff: 1.5-log2-fold change, FDR: 0.001) used for unsupervised clustering. Color legend for BCP-ALL subtypes is the same as in (A). Supplementary Figure S11 and Supplementary Tables S7-S28 provide detailed information on the derived gene sets. **(C)** Canonical signaling pathways (KEGG, HALLMARK gene sets; MSigDB) were used for single sample gene set enrichment analysis using the BCP-ALL subcohort from (B) for balanced representation of all subtypes. Enrichment scores for top variable enriched pathways are shown.

## Discussion

Risk stratification based on molecular disease subtypes has contributed to the remarkable improvement in outcomes of patients with BCP-ALL in the last decades and has provided guidance for target specific treatments. Current nosology of BCP-ALL includes up to 26 specific subtypes (WHO-HAEM5/ICC)^1,2^, exceeding the capability of cytogenetic and molecular genetic techniques which have so far been combined for molecular subtype allocation. Transcriptome sequencing provides informative gene expression profiles and allows identification of underlying driver gene fusions and more recently also driver single nucleotide variants and karyotypes. Analysis of gene expression profiles for molecular subtype allocation is still not standardized, despite its potential for validating genomic driver calls and for subtype allocation of samples with missed genomic drivers.^4^

We have developed ALLCatchR, a pre-trained machine learning classifier which allows molecular subtype allocation in independent hold-out data with >95% accuracy. ALLCatchR is the only tool which systematically provides allocation to all gene expression defined subtypes of the ICC classification, including novel *CDX2/UBTF* ALL^4,26–28^ and *CEBP/ZEB2*^29–31^. Comparable published approaches (ALLSorts, ALLspice) also achieved accurate predictions. However, ALLCatchR achieved superior performance through enabling more correct subtype allocations especially in a real-world adult BCP-ALL data set from a diagnostic laboratory (MLL)^8^, probably due to incorporation of similar data from an independent adult cohort in the training set (GMALL)^3,4^. Immunophenotyping is a routine diagnostic in BCP-ALL and provides putative differentiation stages of origin with ‘pro-B’ immunophenotype used as high-risk marker in some treatment stratification systems. EGIL definitions^20^ were derived from murine B lymphopoiesis. Projecting BCP-ALL samples to our newly established reference of normal lymphopoiesis yielded novel insights into differentiation stages of origin shared between BCP-ALL subtypes. Interestingly, *KMT2A* and *PAX5* P80R ALL, showed a strong proximity to normal pro-B cells, the most immature B lymphoid stage analyzed. These observations are in line with very recent single cell analyses suggesting a pro-B or even pre-pro-B origin of *KMT2A* ALL^24,32^ and murine models of *PAX5* P80R ALL showing that homozygous *PAX5* P80R induces a pro-B differentiation arrest in lymphopoiesis before full transformation through acquisition of additional driver events.^25^ Here, ALLCatchR analysis based on our large aggregated reference cohort confirmed these observations of smaller cohorts^24,32^, preclinical models^25^ and previous assumptions on red-directed *PAX5* functionality in *PAX5* P80R ALL^3,5^. Gene-expression-based definitions of developmental stages in BCP-ALL were more closely related to BCP-ALL subtypes than immunophenotypes, suggesting that selection for leukemogenic drivers occurs in a differentiation-stage specific manner.

ALLCatchR is based on the largest cohort of BCP-ALL gene expression profiles across age groups and molecular subtypes available till date. We make use of this aggregated data to provide subtype defining gene sets for normal and leukemic B lymphopoiesis as an independent research resource. Although only a small minority of samples remain ‘unassigned’, novel subtype candidates are being discussed (e.g.; *IDH1/2* mutated ALL, Low hyperdiploid ALL)^5,26^. ALLCatchR is a freely available open-source tool providing a conceptual and technical framework which can easily be extended for incorporation of novel subtypes and additional predictive models. When combined with already establishes approaches for calling of genomic drivers (e.g., gene fusions), ALLCatchR will complement the essential prerequisites for the transition of RNA-Seq from research to routine application in clinical diagnostics.

## Acknowledgements

This study was in part funded by the Deutsche Forschungsgemeinschaft (DFG, German Research Foundation) – project number 444949889 (KFO 5010/1 Clinical Research Unit ‘CATCH ALL’ to L.B., A.H., M.P.H., M.N., M.B., and C.D.B.), and project number 413490537 (Clinician Scientist Program in Evolutionary Medicine to B.T.H.) and Deutsche Jose Carreras Leukämie Stiftung (DJCLS 01R/2016 to L.B. and C.D.B, DJCLS R 15/11 and DJCLS 06R/2019 to M.Br.) and the Czech Health Research Council (NU20-07-00322 to M.Z. and J.T.)

We gratefully appreciate critical contributions from Saskia Kohlscheen and Matthias Ritgen for the development of the healthy donor FACS sort panel and Monika Szczepanowski for contributing to sample collection critical discussion of the manuscript. We are indebted to Christian Peters and Esther Schiminsky for performing the FACS sorts.

## Author contributions

T.B., M.Br., C.D.B. and L.Bas. designed the study; T.B. and L.Bas. established models for molecular subtype allocation and B cell developmental stages and developed the classifier; B.T.H., L.Bas. and C.D.B. conceived the clinical trial to obtain healthy bone marrow samples; B.T.H., E.A. and L.Bas. established the normal donor FACS panel, B.T.H. and M.Bu. performed FACS sorting; T.B., B.T.H, A.M.H., N.K., L.Bar., S.B., J.K., M.B. established bioinformatic workflows and performed analyses of BCP-ALL and healthy donor gene expression profiles; J.Z. and Chr. K. developed and tested the CRAN package for ALLCatchR distribution, W.W., M.Z., Z.A., P.C., G.C., M.S., M.N., N.G., A.K.B., J.T., C.H. contributed BCP-ALL sequencing data and validated ground truth and/or contributed to the classifier concept; L.Bas. and C.D.B. supervised the project; T.B., C.D.B. and L.Bas. drafted the first version of the manuscript; all authors revised and approved the final version of the manuscript.

## Competing Interests

The authors have no competing interests to declare.

## Data Availability Statement

ALLCatchR is freely available as an R-package through https://github.com/ThomasBeder/ALLCatchR. Transcriptome sequencing data of bone marrow samples from healthy donors were deposited at the European Genome Phenome archive. The accession number will be provided after acceptance of the manuscript. BCP-ALL transcriptome profiles haven been deposited in open or controlled access archives by the authors of the original publications.

## Notes

### Competing Interest Statement

The authors have declared no competing interest.

