## Supplementary Figures for "The gene expression classifier ALLCatchR identifies B-precursor ALL subtypes and underlying developmental trajectories across age"

### Supplementary Figure 1

#### ALLCatchR

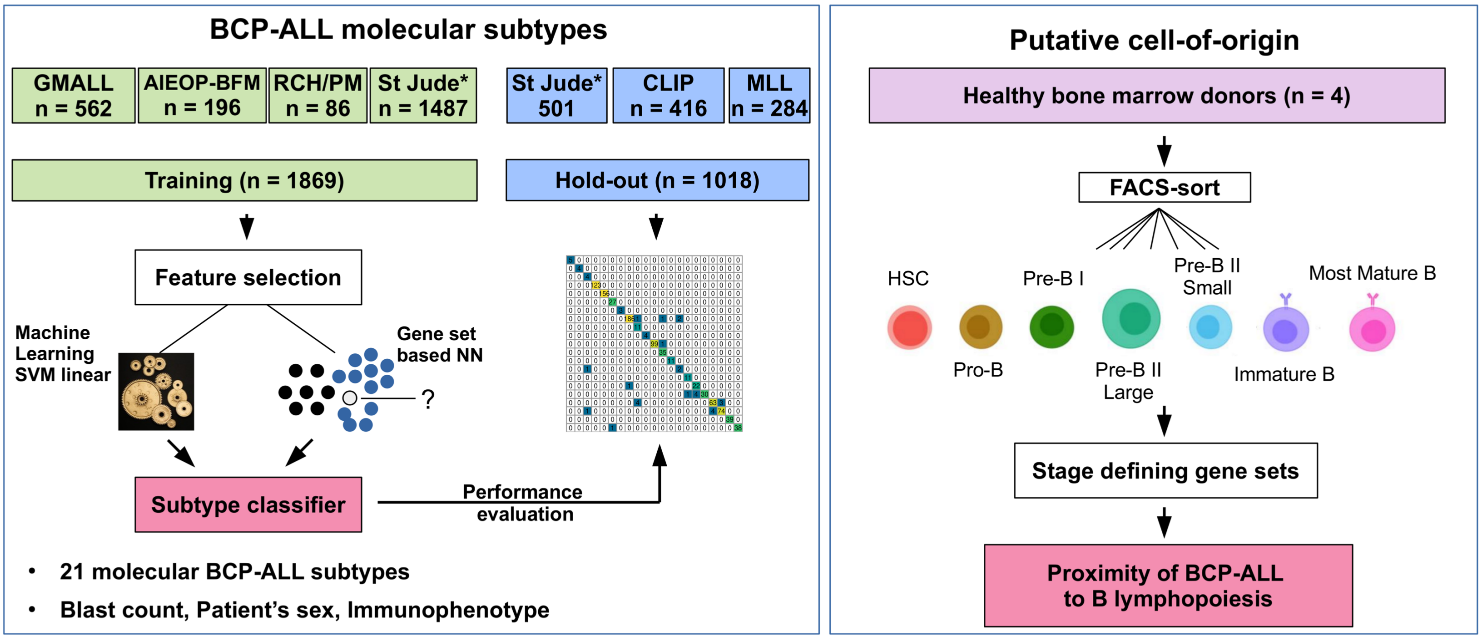


*data from n=10 sub-cohorts, split into training (n=9) and hold-out (n=1)

**Supplementary Figure 1. ALLCatchR workflow for BCP-ALL molecular subtype classification and prediction of the putative cell of origin.** Gene wise count data of a total of n=3,532 BCP-ALL patients from RNA-Seq experiments comprising six datasets were included in this study. Four data sets (n=1869) were used for training and validation was performed on three hold-out data sets (n=1018). ALLCatchR is a compound classifier based on deterministic linear SVM predictions and a classifier based on sample-to-sample distances to subtype specific gene sets.

### Supplementary Figure 2

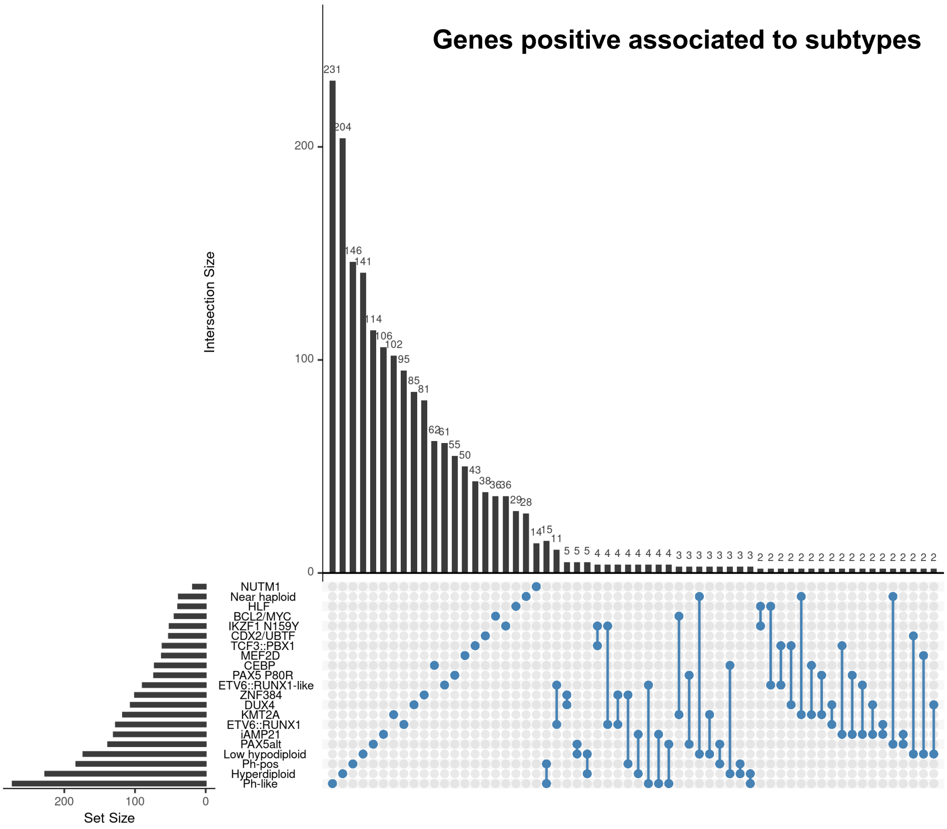


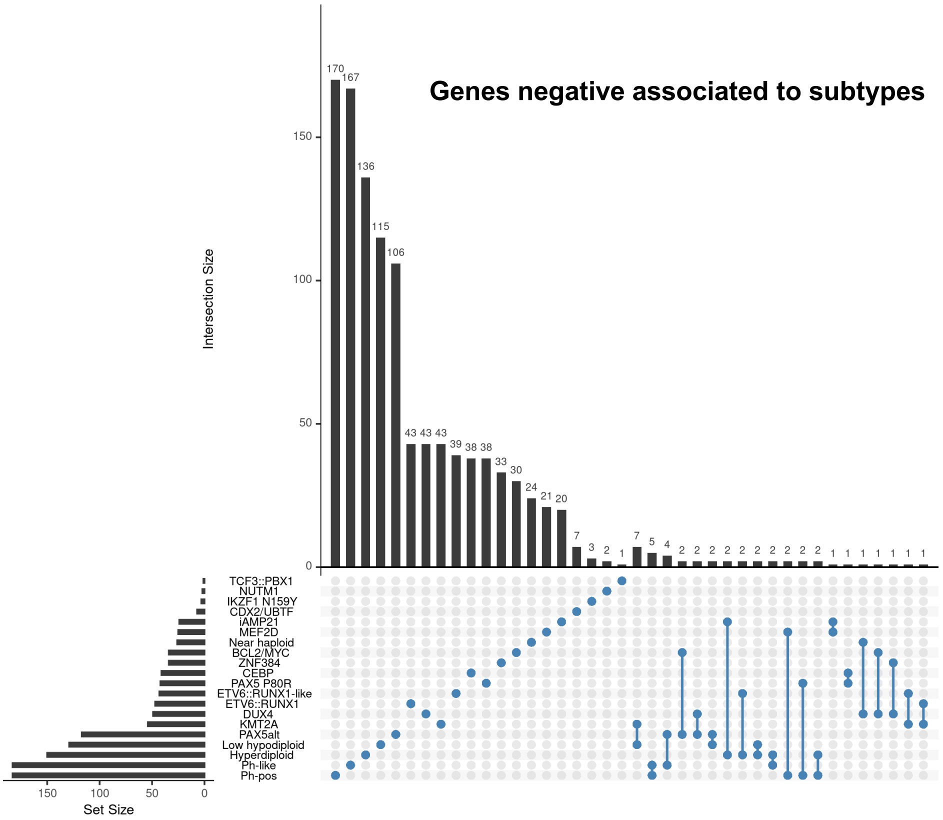


**Supplementary Figure 2. The 2,802 LASSO genes used for training ALLCatchR are highly specific for BCP-ALL molecular subtypes.** Upset plots showing the number of genes (Set Size) for each subtype and the number of unique and shared genes between subtypes (Intersection Size).

### Supplementary Figure 3

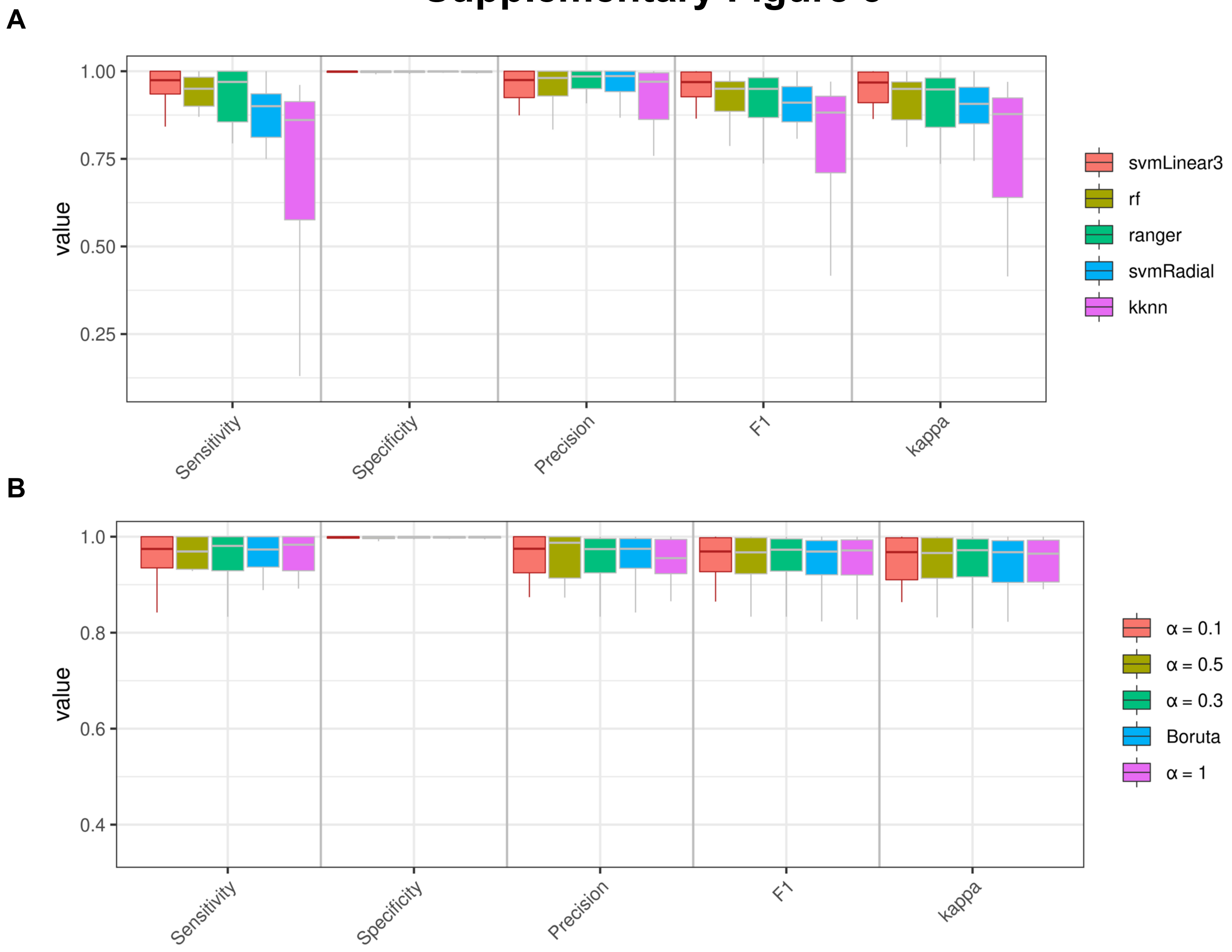


**Supplementary Figure 3. Choice of machine learning algorithm contributes to subtype prediction performance. (A)** Performance of different machine learning algorithms was tested on a combined set of training and test data (n=2,887) with genomic defined subtype allocations. Box plots showing classification performance of different machine learning algorithms[12] (svmLinear3, rf, ranger svmRadial and kknn) for averages of individual subtypes. Linear SVM was selected for ALLCatchR because of superior performance on the training data out of five methods tested with an overall accuracy of 0.963 followed by radial SVM (accuracy: 0.957) and KNN (accuracy: 0.901). Overall accuracies are skewed for the impact of large subtypes. Linear SVM was still the best performing method when all subtypes were weighted equally with an average sensitivity of 0.950±0.085 and specificity of 0.998±0.003 **(B)** Features selection methods LASSO[12] and Boruta[15] were applied to account for linear feature to class label interactions as well as non-linear interactions. Different alpha parameters were set in LASSO, where higher values result in a more stringent selection of features. This resulted in different numbers of selected genes for machine training ranging from n=2,802 (LASSO α=0.1), to n=973 (LASSO α=1). The comparison showed that linear SVM performed stable independent of the feature selection used.

### Supplementary Figure 4

**A**

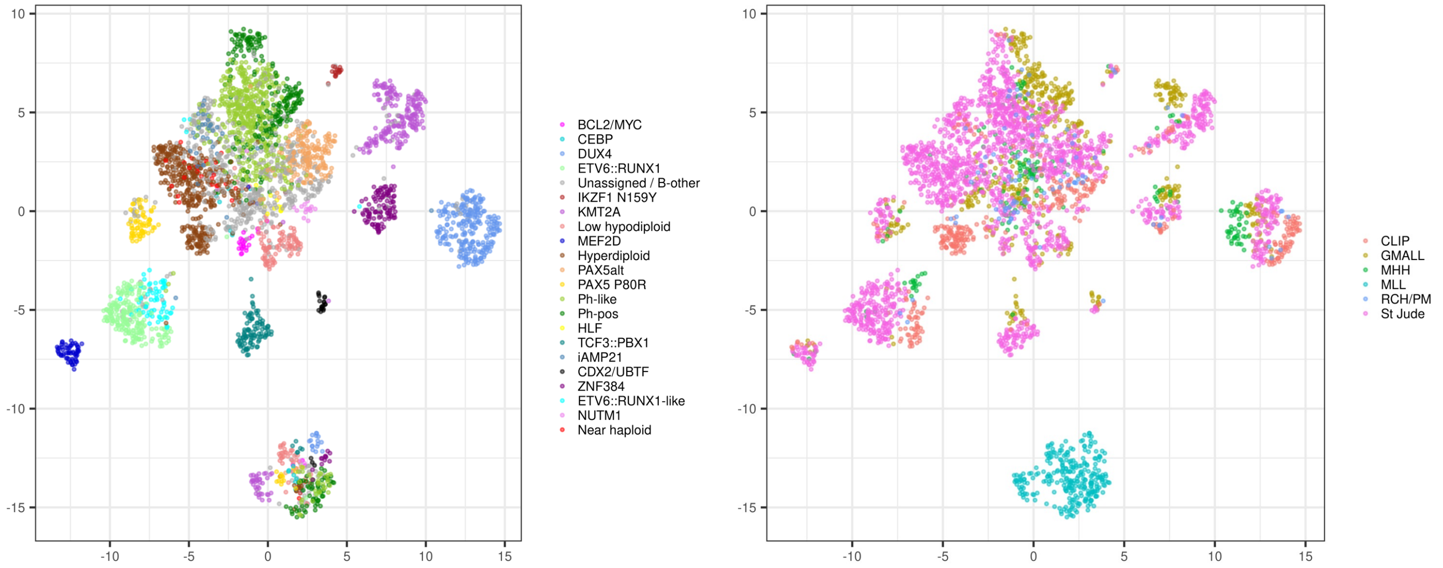


**B**

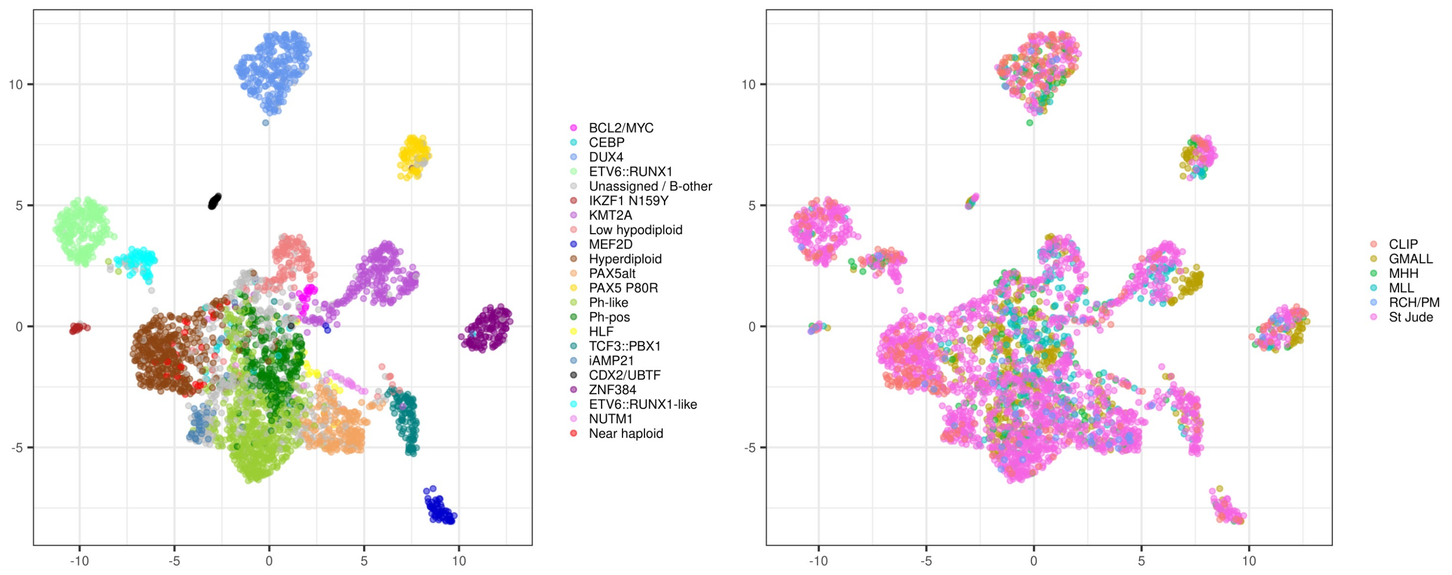


**Supplementary Figure 4. Samples separate according to BCP-ALL molecular subtypes based on LASSO genes. (A)** UMAP plots depict all samples based on expression of 2,802 LASSO genes (Supplementary Table S3) used for training ALLCatchR. Samples group according to their subtypes, but batch effects remain (e.g; MLL cohort). **(B)** Using Lasso gene sets used in (A), single sample gene set analysis was performed with singscore for the n=1,869 training samples, which allowed the computation of a total enrichment score for each sample to each subtype specific gene set. These 21 total enrichment scores for each sample can be used to clearly separate the subtypes as shown here. No more cohort-specific batch effects are observed.

### Supplementary Figure 5

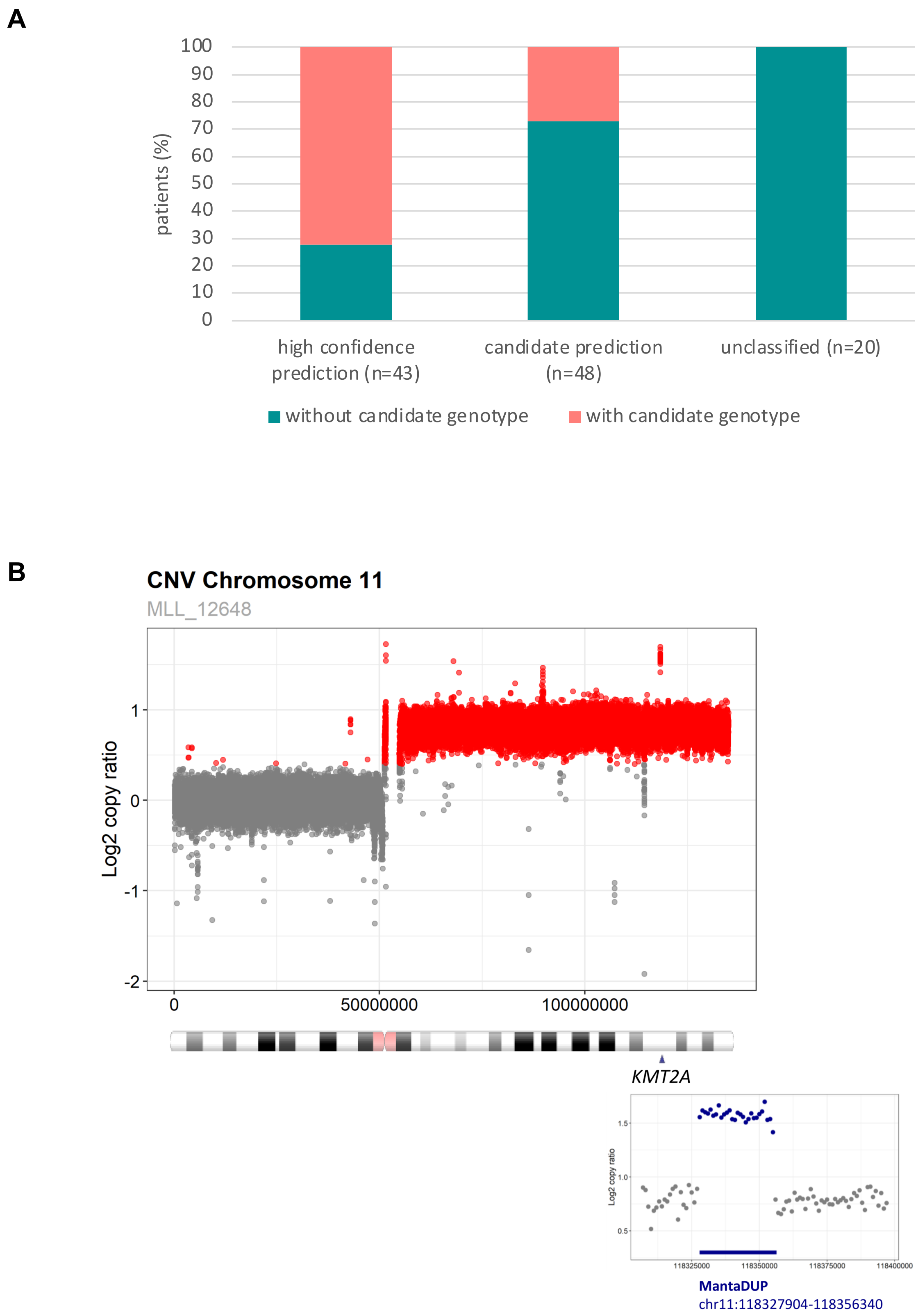


**Supplementary Figure 5. Identification of novel candidate subtype assignments in ’unassigned / B-other’ samples. (A)** A total of n=111 samples of the hold-out-studies were defined ‘unassigned / B-other’ in the original studies. ALLCatchR provided subtype allocations for n=91 of these samples (Supplementary Table S4). The proportions of samples where these predictions could be confirmed be corresponding genomic aberrations are shown here. **(B)** In one sample with a high-confidence KMT2A prediction but absence of KMT2A fusion an i(11)(q10) was identified by whole genome sequencing with and additional intrachromosomal tandem duplication at breakpoint chr11:118327904-118356340.

### Supplementary Figure 6

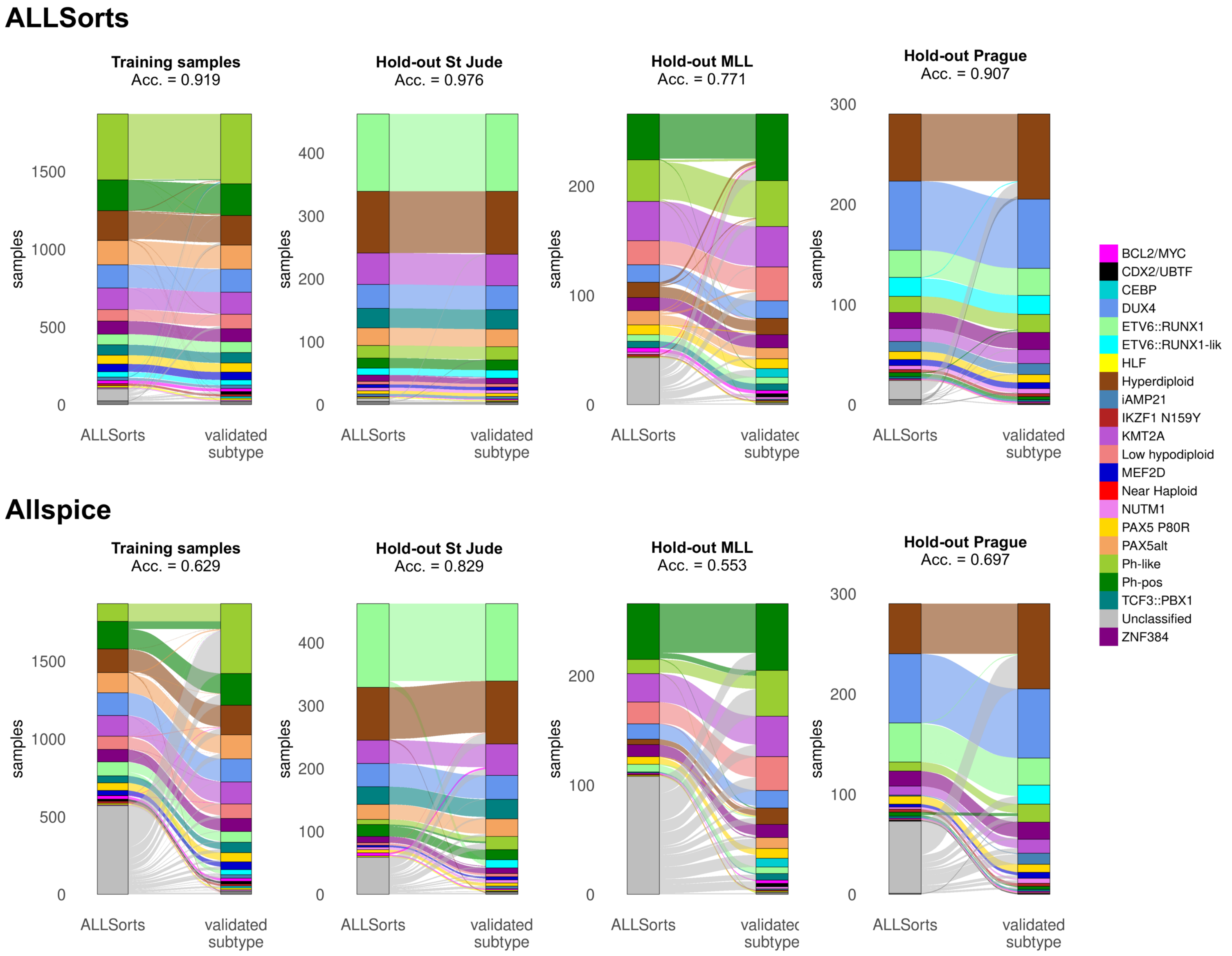


**Supplementary Figure 6. Sankey diagrams showing the subtype classification performance of ALLSorts and Allspice on the training and hold-out data sets.**

### Supplementary Figure 7

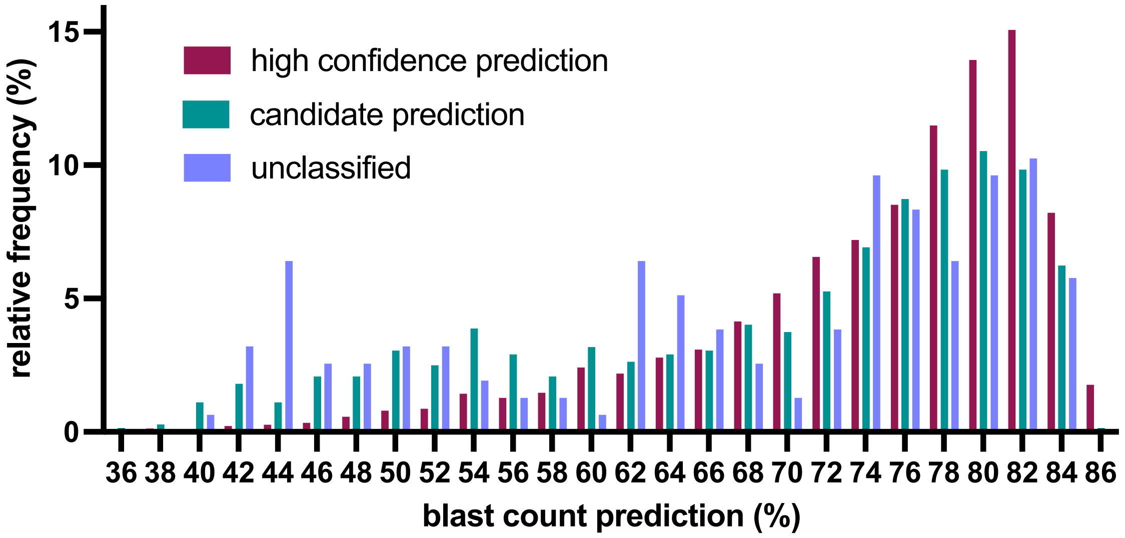


**Supplementary Figure 7. Predicted blast counts according to BCP-ALL subtype allocations performed by ALLCatchR.** Blast counts were computed for all samples of the combined cohort based on training as described in Figure 3 of the manuscript. The relative frequency distribution of blast counts is depicted here in relation to the quality of ALLCatchR BCP-ALL subtype allocations. Samples with low blast count estimates are enriched for 'candidate' or 'unclassified' predictions.

### Supplementary Figure 8

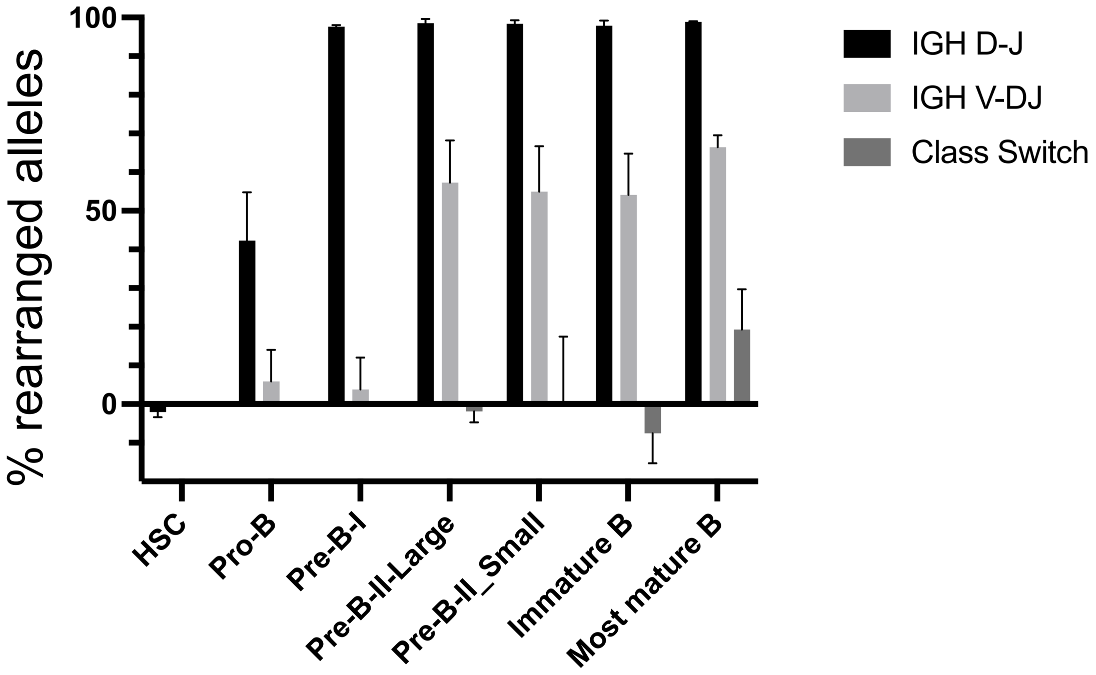


**Supplementary Figure 8. Analysis of immunoglobulin rearrangements using droplet PCR.** The fraction of rearranged IGH-alleles was measured using a ddPCR that targets the sequences that are deleted during the D-J-, V-DJ- and class switch rearrangements, respectively, together with DNM3 as a reference gene in a multiplex ddPCR setup. The PCR program and the primers and probes were used as previously described. (1) The probes targeting the respective IGH sequences were labeled with a FAM fluorophore and the DNM3 probe with a HEX fluorophore. The 20 µL PCR mixture contained 1X ddPCR Supermix for Probes (no dUTPs) (Bio-Rad), 300 nM of each primer, 100 nM of each probe and 50 ng DNA of interest. Droplet generation and readout was performed as previously specified. (2) The percentage of rearranged alleles was calculated as described. (1)

References to Supplementary Figure S8:

1) Zoutman WH, Nell RJ, Versluis M, et al. A novel digital PCR-based method to quantify (switched) B cells reveals the extent of allelic involvement in different recombination processes in the IGH locus. Mol. Immunol. 2022;145:109–123.

2) Zoutman WH, Nell RJ, van der Velden PA. Usage of Droplet Digital PCR (ddPCR) Assays for T Cell Quantification in Cancer. 2019;1–14

### Supplementary Figure 9

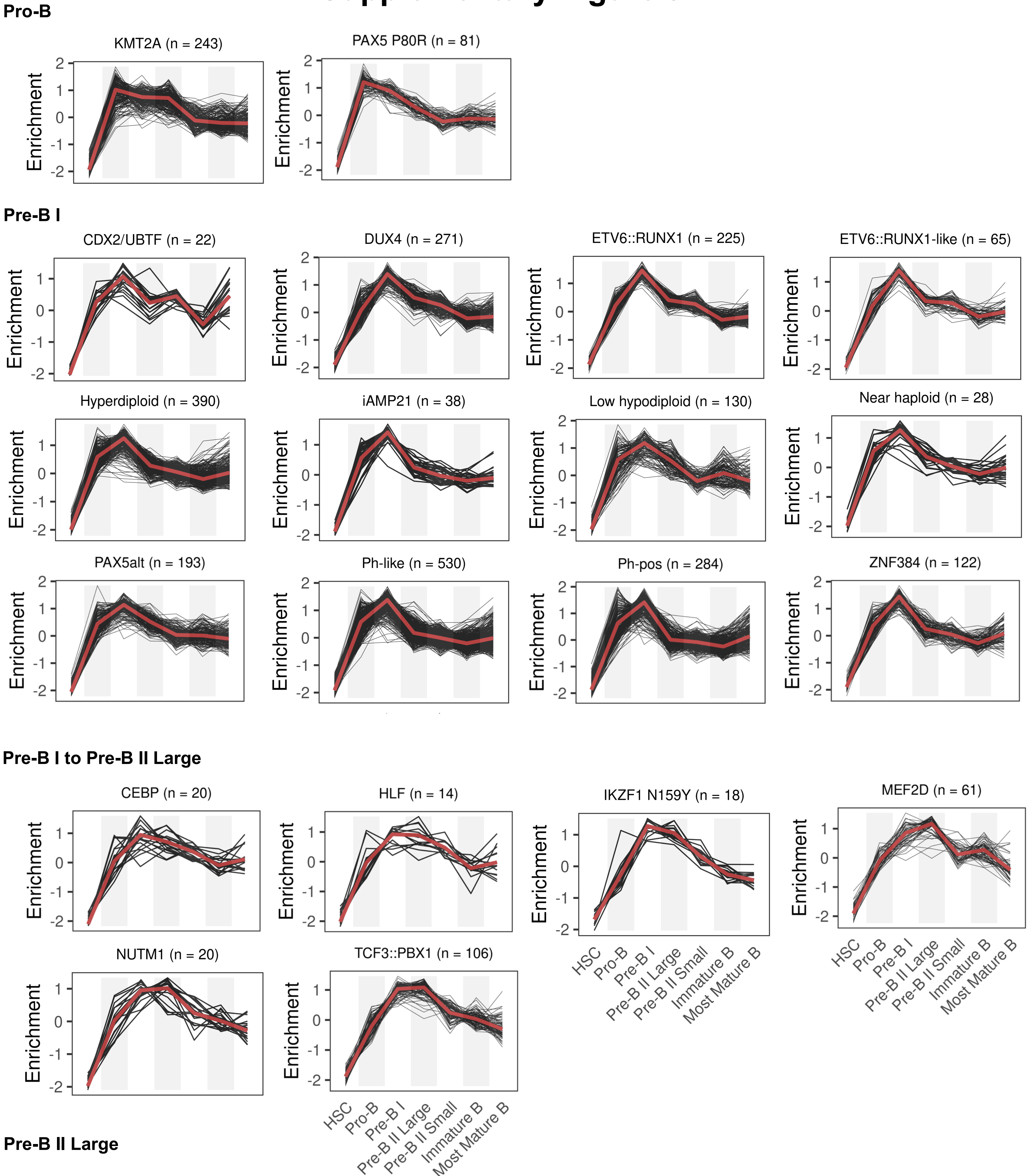


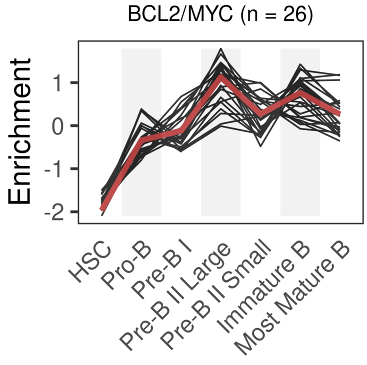


**Supplementary Figure 9. Gene set enrichment analysis of BCP-ALL samples shows distinct pattern shared between BCP-ALL subtypes.** Single sample gene set enrichment analysis was performed for BCP-ALL samples with defined molecular subtype (n=2,887) and lymphopoiesis-stage specific gene sets (Figure 4, main manuscript). Enrichment patterns shared between BCP-ALL subtypes suggest common B lymphopoiesis stages of origin. Condensed heatmap presentations of this data is shown in Figure 4D and Supplementary Figure S10. KMT2A and PAX5 P80R individual enrichment profiles are shown in Figure 4E of the main manuscript and are included here for complete overview.

### Supplementary Figure 10

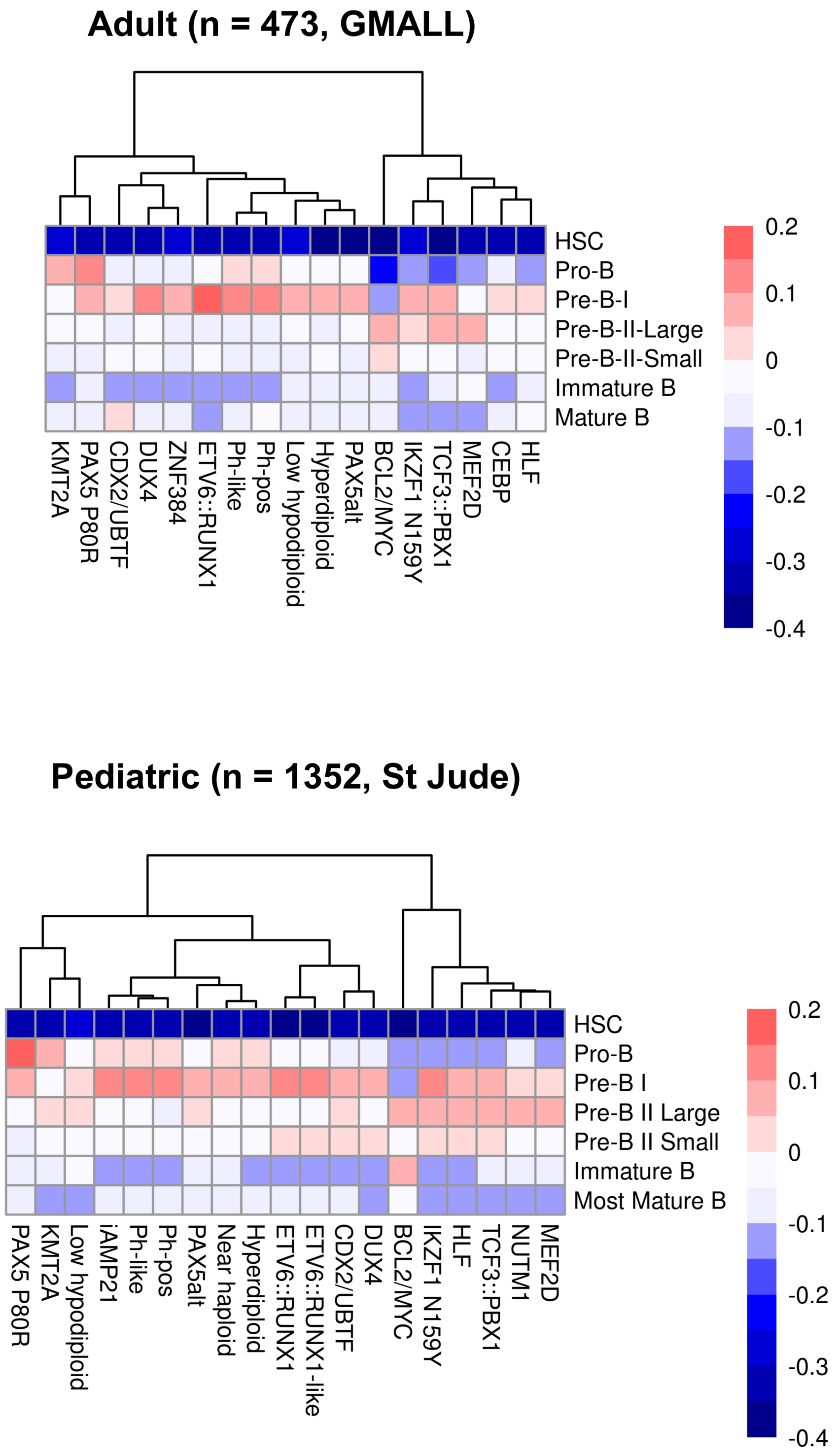


**Supplementary Figure 10. Consistent associations to B progenitor gene sets were found in adults and children.** Hierarchical clustering of subtypes based on gene expression signatures specific to the seven B lymphopoietic differentiation stages. Adult samples from GMALL and pediatric of St Jude are shown and enrichment scores were averaged for the individual subtypes.

### Supplementary Figure 11

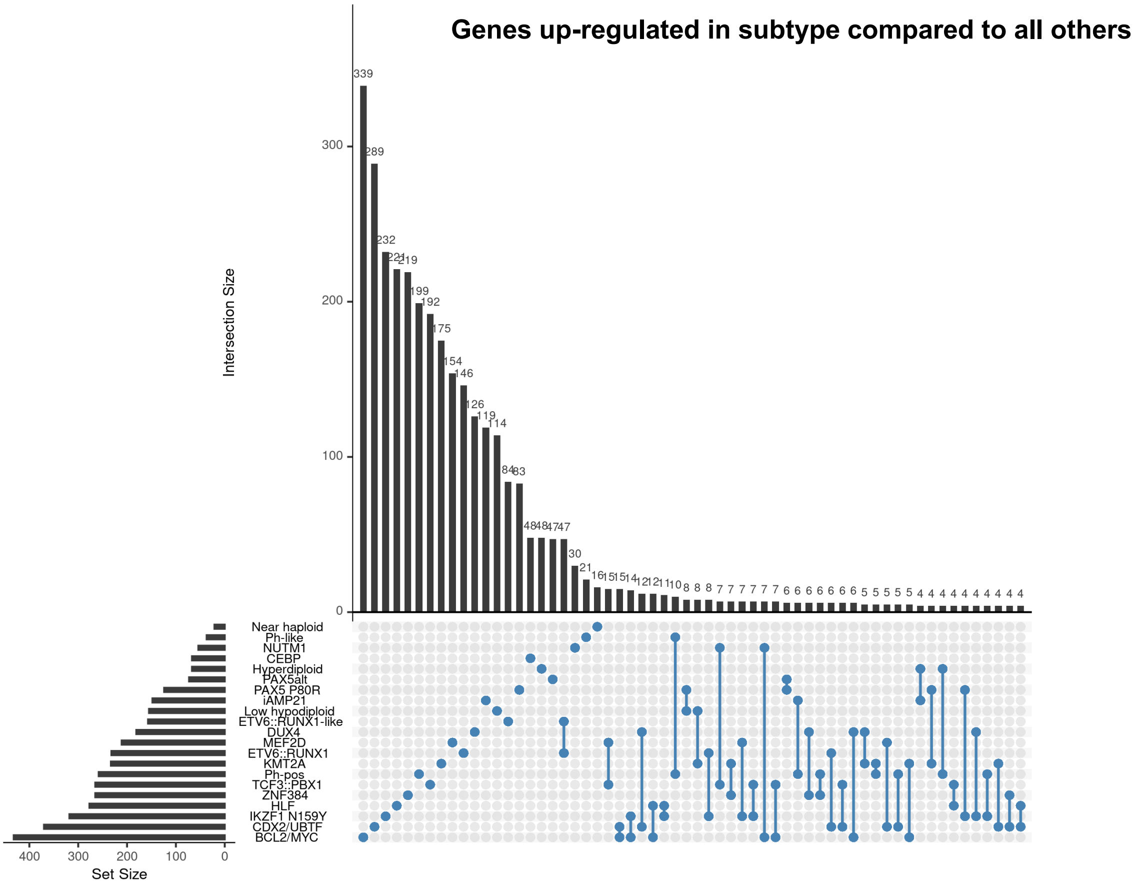


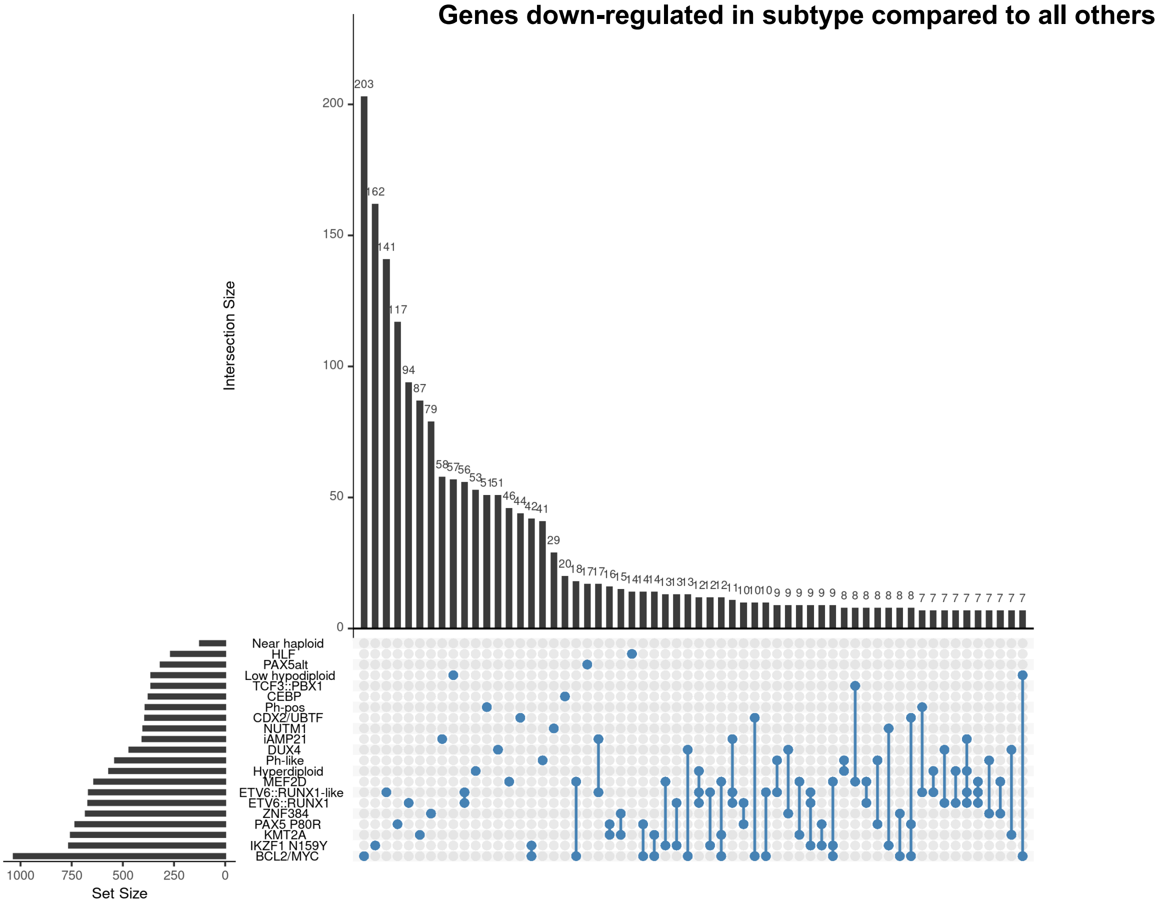


**Supplementary Figure 11. Differential expressed genes between BCP-ALL molecular subtypes.** Upset plots showing the number of genes (Set Size) for each subtype and the number of unique and shared genes between subtypes (Intersection Size).
